# Neuroplasticity of speech-in-noise processing in older adults assessed by functional near-infrared spectroscopy (fNIRS)

**DOI:** 10.1101/2023.09.23.559144

**Authors:** Guangting Mai, Zhizhao Jiang, Xinran Wang, Ilias Tachtsidis, Peter Howell

## Abstract

Functional near-infrared spectroscopy (fNIRS), a non-invasive optical neuroimaging technique that is portable and acoustically silent, has become a promising tool for evaluating auditory brain functions in hearing- vulnerable individuals. This study, for the first time, used fNIRS to evaluate neuroplasticity of speech-in-noise processing in older adults. Ten older adults, most of whom had moderate-to-mild hearing loss, participated in a 4-week speech-in-noise training. Their speech-in-noise performances and fNIRS brain responses to speech (auditory sentences in noise), non-speech (spectrally-rotated speech in noise) and visual (flashing chequerboards) stimuli were evaluated pre- (T0) and post-training (immediately after training, T1; and after a 4-week retention, T2). Behaviourally, speech-in-noise performances were improved after retention (T2 vs. T0) but not immediately after training (T1 vs. T0). Neurally, we intriguingly found brain responses to speech vs. non-speech decreased significantly in the left auditory cortex after retention (T2 vs. T0 and T2 vs. T1) for which we interpret as suppressed processing of background noise during speech listening alongside the significant behavioural improvements. Meanwhile, functional connectivity within and between multiple regions of temporal, parietal and frontal lobes was significantly enhanced in the speech condition after retention (T2 vs. T0). We also found neural changes before the emergence significant behavioural improvements. Compared to pre-training, responses to speech vs. non-speech in the left frontal/prefrontal cortex were decreased significantly both immediately after training (T1 vs. T0) and retention (T2 vs. T0), reflecting possible alleviation of listening efforts. Finally, connectivity was significantly decreased between auditory and higher-level non-auditory (parietal and frontal) cortices in response to visual stimuli immediately after training (T1 vs. T0), indicating decreased cross-modal takeover of speech-related regions during visual processing. The results thus showed that neuroplasticity can be observed not only at the same time, but also *before* behavioural changes in speech-in- noise perception. To our knowledge, this is the first fNIRS study to evaluate speech-based auditory neuroplasticity in older adults. It thus provides important implications for current research by illustrating the promises of detecting neuroplasticity using fNIRS in hearing-vulnerable individuals.

## 1 Introduction

How the brain processes speech is an important topic in auditory cognitive neuroscience. A long-standing focus is to study the brain functions in hearing-vulnerable populations such as older adults and hearing-impaired listeners who experience challenges in speech and language perception (see reviews: Peelle and Wingfield, 2016; Slade et al., 2020). This current study asks questions on how contemporary sophisticated functional neuroimaging techniques can help us practically study this essential topic. Over the years, studies have used techniques, such as functional magnetic resonance imaging (fMRI) and positron emission tomography (PET), to illustrate the breakdown of brain processing of speech and language in older and hearing-impaired listeners (Wong et al., 1999; Wong et al., 2009; Peelle et al., 2011; Vaden et al., 2015, 2016; Vogelzang et al., 2021). Both fMRI and PET detects dynamics of cerebral haemoglobin (haemodynamic responses) at different regions of the brain capturing neural responses with high special resolution and has been used in auditory research (e.g., Zatorre et al., 1996; Zatorre, 2001; Hall et al., 1999, 2009; Peelle, 2014). Using these techniques, studies have observed altered neural sensitivity to speech signals in the auditory cortices (Wong et al., 1999; Wong et al., 2009; Peelle et al., 2011) as well as abnormal neural responses at higher-level non-auditory, cognitive regions in these individuals compared to normal-hearing young adult listeners (Wong et al., 1999; Wong et al., 2009; Vaden et al., 2015, 2016; Vogelzang et al., 2021). While both fMRI and PET have been widely used, they also face limitations in auditory research. For example, both techniques can be expensive and may not be always easy to use for large-scale studies in clinical populations (Boas et al., 2014; Pinti et al., 2020). Also, fMRI generates loud extraneous scanning noise that can cause problems for assessing auditory functions (Scarff et al., 2004; Gaab et al., 2007). Furthermore, hearing protheses, like hearing aids and cochlear implants, can have intensive magnetic interference with MRI scanning (Saliba et al., 2016; Basura et al., 2018; Harrison et al., 2021) such that hearing aid and cochlear implant users are largely excluded from fMRI research. For PET, although it is noise-free and does not have magnetic interactions with hearing protheses, it requires an invasive procedure, i.e., injection of radioactive isotopes (Johnsrude et al., 2002), making it unsuitable for repetitive use in clinical populations. Besides fMRI and PET, functional near-infrared spectroscopy (fNIRS) is another promising technique to study the neural processes of auditory and speech perception (Pollonini et al., 2014; Wiggins et al., 2016; Defenderfer et al., 2017, 2021; Wijayasiri et al., 2017; Lawrence et al., 2018; Mushtaq et al., 2021; Zhou et al., 2022). fNIRS is an optical imaging technique that illuminates scalp of the brain using near-infrared light and measures the intensity of light returning from cortical areas through which haemodynamic responses are estimated (Boas et al., 2014; Pinti et al., 2020). Nowadays it has become more advantageous and practical to use in hearing-vulnerable populations to study their auditory brain functions (Boas et al., 2014; Pinti et al., 2020). Here, we focus on how fNIRS may be feasible and show promises of measuring changes in neural processing of speech in hearing-vulnerable populations.

Compared to fMRI or PET, fNIRS is more portable and relatively less expensive hence easier to use in laboratory environments for clinical populations (Boas et al., 2014; Pinti et al., 2020). Also, compared to fMRI, fNIRS is acoustically silent which is crucial for auditory experiments in those who face challenges in hearing and speech. Furthermore, unlike PET, fNIRS is non-invasive, making it more suitable for repeated measurements, e.g., in longitudinal studies, for clinical populations (Saliba et al., 2016; Basura et al., 2018; Harrison et al., 2021). Lastly, fNIRS is compatible with people who wear hearing protheses like hearing aids and cochlear implants which can have intensive magnetic interference with MRI scanning (Saliba et al., 2016; Basura et al., 2018; Harrison et al., 2021). Recent research has successfully used fNIRS to illustrate the neural processes of hearing and speech perception in hearing-vulnerable populations. For instance, using fNIRS, Olds et al. (2016) showed that cochlear implant patients with good speech perception exhibited greater auditory cortical activations in response to intelligible than unintelligible speech whilst those with poor perception did not show distinguishable activations, revealing the association between speech perception and cortical activities. Previous studies have also shown successes in detecting listening efforts using fNIRS in older and hearing-impaired listeners. Rovetti et al., (2019) showed that reduction of fNIRS prefrontal cortical activations (reflecting alleviation in listening effort) during an auditory N-back task is associated with the use of hearing aids in older adults with hearing loss. Sherafati et al. (2022) showed greater fNIRS prefrontal cortical activations in cochlear implant patients than normal-hearing controls during spoken word listening tasks, reflecting greater listening efforts in the implanted patients. fNIRS also demonstrated promises in detecting cross-modal activations in relation to speech perception in the hearing-impaired. For instance, Anderson et al. (2017) showed that better speech perception in cochlear implant patients is associated with enhanced fNIRS cross-modal activations (auditory cortical responses to visual speech). Fullerton et al. (2022) further showed better speech perception is associated with functional connectivity between auditory and visual cortices in response to visual speech in implanted patients.

Despite these successes of the use of fNIRS and its unique advantages, previous research also confronted limitations of this technique. For example, compared to neuroelectromagnetic methods like electroencephalography (EEG) and magnetoencephalography (MEG), fNIRS measures haemodynamic responses that are sluggish, so it is unable to capture fine-grained timing information of the neural signals (Pinti et al., 2020). Also, its restricted depth of optode penetration makes it only detects neural activations occurred in the outer cortices with a relatively sparse spatial resolution compared to fMRI and PET which can further capture activities within sulci and deep into medial cortices (Pinti et al., 2020). Hence, it is worth noticing these limitations due to which some brain functions may not be easily detected through fNIRS. Therefore, evaluating the feasibility of this technique as discussed above is an important step to confirm its great promises in auditory research. However, most of these efforts so far have focused on cross-sectional experiments and it is unclear how *changes* in brain functions over time could be feasibly detected by fNIRS. Such changes are referred as ‘neuroplasticity’, which reflects the capacity of the brain to undergo functional reorganization across time (Innocenti, 2022). Observing this plasticity is important because it should pave the way for future research into the neural mechanisms underlying the behavioural changes, especially in older adults and hearing-impaired populations who have shown the potential to improve their speech perception after proper speech-based training interventions (Stropahl et al., 2020; Bieber and Gordon-Salant, 2021). Clinically, it can help identify those who have strong potentials for positive neuroplastic changes so that individualized treatments can be properly designed (Cramer et al., 2011; Nahum et al., 2013).

The current study aimed to assess the promises of using fNIRS to detect auditory neuroplasticity through a longitudinal experiment in older adults, most of whom had mild-to-moderate hearing loss. Participants received a 4-week home-based speech-in-noise training and their brain activities were measured by fNIRS over the speech- and language-related cortical areas (temporal, parietal and frontal regions, see Poeppel and Hickock, 2007) both before and after training. Neural responses at various brain regions of interest as well as functional connectivity were examined during an auditory and a visual test and were compared between sessions before and after training. In the auditory test, participants listened to speech (spoken sentences) and non-speech stimuli (spectrally-rotated versions of speech controlling for acoustic complexity to examine speech specificity) presented in noisy backgrounds. We expect increased auditory cortical activities reflecting greater auditory sensitivity after training as well as decreased left frontal/prefrontal cortical activities reflecting reduced listening efforts (Wild et al., 2012; Wijayasiri et al., 2017; Rovetti et al., 2019; Sherafati et al., 2022). We also expect enhancements in brain connectivity reflecting better coordination between language-related areas (Poeppel and Hickock, 2007). In the visual test, participants were exposed to speech-unrelated visual stimuli (flashing chequerboards). Previous research has reported that such stimuli can elicit greater auditory cortical activities in hearing-impaired people reflecting cross-modal maladaptation associated with poorer speech perception (Campbell and Sharma, 2014; Chen et al., 2015; Corina et al., 2017). We expect that this maladaptation would be reduced after training (i.e., reduced auditory cortical activities and/or reduced connectivity between auditory cortex and higher-order parietal and frontal speech-related areas in response to visual stimuli). We anticipate that the observed longitudinal changes should provide new insights into possible underlying mechanisms for changes in speech-in-noise perception over time.

## 2 Methods and Materials

This study was approved by the UCL Research Ethics Committee. All participants were consent and reimbursed for their participation.

### 2.1 Participants

Ten right-handed, healthy adult participants (two males) aged between 63 and 78 years (mean = 70, SD = 4.5) were recruited. They were all native British English speakers with no reported histories of neurological, cognitive or language disorders. Their pure-tone audiograms (PTAs) were measured for each ear before the speech-in-noise training using a MAICO MA41 Audiometer (MAICO Diagnostics, Germany) at 0.25, 0.5, 1, 2, 3, 4, 6 and 8 kHz. Two participants had normal hearing (≤ 25 dB HL) at all frequencies ≤ 6 kHz in both ears. The other eight showed a general pattern of mild-to-moderate hearing loss (30–60 dB HL) especially at high frequencies (> 2 kHz) (see **Figure 1**). This therefore matches the real-life scenario where majority of healthy ageing populations suffer from high-frequency mild-to-moderate hearing loss (Gopinath et al., 2009; Humes et al., 2010).

**Figure 1.**
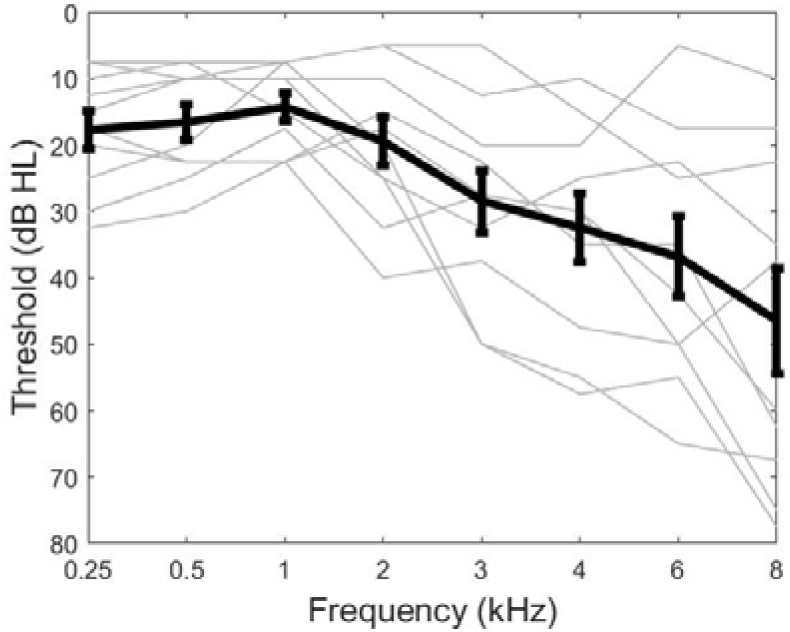
Audiograms of participants averaged across the two ears. Grey lines show the thresholds of individual participants and the black line show the thresholds averaged across participants. Error bars indicate the standard errors of the means.

### 2.2 Design

Participants received a home-based speech-in-noise training through a participant-/patient-friendly App developed by Green et al., (2019). With proper instructions, participants were able to complete the training process by themselves via controlling the Matlab Graphical User Interfaces using a computer tablet at their own home. Training data were saved in an online UCL Research Dropbox in a daily basis so that experimenters could make sure the training was gone through smoothly. During the training, participants listened to storybooks (in British English) spoken by a male and a female speaker sentence-by-sentence presented in background noise and they were asked to identify words within each sentence through multiple-choice word tasks. The background noises were multiple-talker babbles (4, 8 and 16 talker-babbles presented throughout the training in intermixed orders; half males and half females). An adaptive procedure was adopted where the signal-to-noise ratio (SNR) increased/decreased following the decreases/increases in participants’ accuracies over time to keep their attention. The training lasted for 4 weeks, 6 days per week, ∼30 minutes per day.

Their speech-in-noise performances and brain responses to auditory and visual stimuli were measured both before (a day or two before the training as the baseline, T0) and after training (the next day after the training ended, T1; and after an additional 4-week retention period, T2). **Figure 2A** illustrates the study procedure.

**Figure 2.**
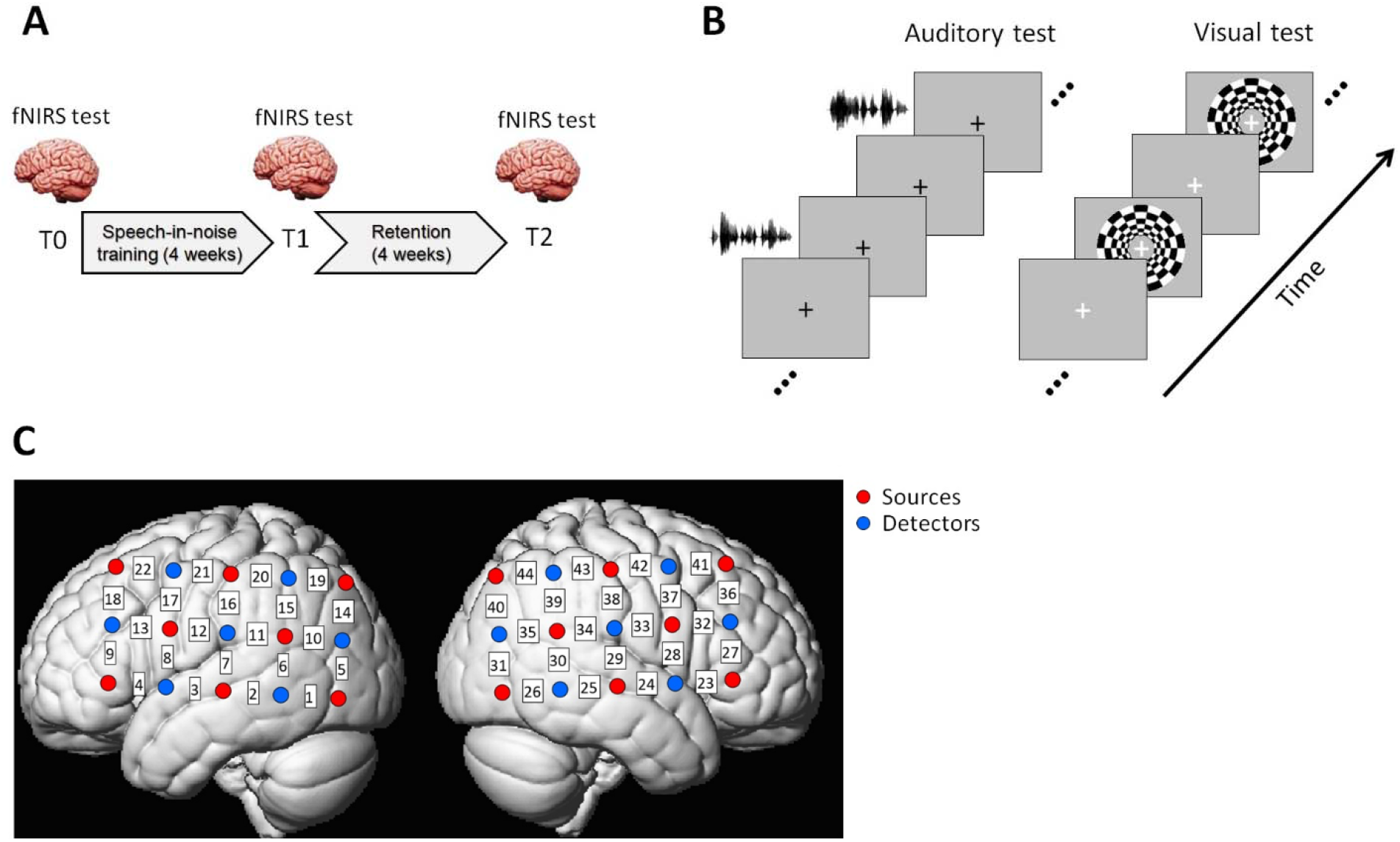
Experiment design. **(A)** Participants completed a 4-week home-based speech-in-noise training and their brain functions were measured by fNIRS before (T0) and after the training (T1 and T2). **(B)** The fNIRS experiment included an auditory test where participants listened to auditory sentences (speech and spectrally rotated speech) and a visual test where participants watched a flashing chequerboard. A block design was adopted with resting blocks interleaved between the auditory/visual stimuli. **(C)** Optode configuration of the fNIRS experiment was two 5-by-3 probe sets that formed 44 channels (22 channels in each hemisphere) over speech- and language-related temporal, parietal and frontal cortical regions (left: left hemisphere; right: right hemisphere). Red and blue circles denote the sources and detectors, respectively. The channel indices were indicated in the white squares between the sources and detectors.

### 2.3 Speech-in-noise tasks

The speech-in-noise performances were measured as participants’ speech reception thresholds (SRT) when they listened to short sentences in noisy backgrounds. The sentences were chosen from the Adaptive Sentence List (ASL), each of which consists of three key (content) words (e.g., ‘He wiped the table’ with key words ‘he’, ‘wiped’ and ‘table’) spoken by a male native British English speaker (MacLeod and Summerfield, 1990). Participants were seated in a quiet room listening to 30 sentences under an 8-talker babble noise (the same 8- talker babbles as in the training) via inserted earphones (ER-3 insert earphone, Intelligent Hearing Systems, USA). They were required to verbally report as many words as they could for each sentence. The signal-to-noise ratio (SNR) was initially set at 6 dB for the first sentence (for which all participants were able to recognize all key words) and was decreased by 4 dB for subsequent sentences until < 50% words (i.e., < 2 words) were correctly reported. SNR was then increased/decreased by 2 dB when word correctness was smaller/greater than 50% for each of the following sentences. The SRT was measured as the mean SNR across all reversals at the step size of 2 dB (Schoof and Rosen, 2014). Therefore, lower SRT reflects better speech-in-noise performance. The overall sound level (sentence plus noise) was calibrated and fixed at 75 dB SL. The procedure was controlled using Matlab 2016a (Mathworks, USA) with key words for each sentence appearing on the computer screen seen only by the experimenters. The ‘loose keyword scoring’ approach was followed, meaning that a reported word was considered correct as long as it matched the root of a key word (e.g., ‘stand’ was considered correct for the keyword ‘stood’) (Macleod and Summerfield, 1990). There were 6 practice sentences prior to each formal test.

### 2.4 fNIRS experiment

#### 2.4.1 Optode placements

Brain haemodynamic responses were recorded by a continuous-wave fNIRS system (ETG-4000, Hitachi Medical, Japan; sample rate of 10 Hz) that uses two wavelengths of light at 695 and 830 nm to allow the estimates of changes in both oxy- (HbO) and deoxy-haemoglobin (HbR). The haemodynamics were measured using two 5-by-3 optode probe sets (8 sources and 7 detectors with a fixed source-detector distance of 3 cm on both hemispheres), hence 44 channels covering much of the temporal, parietal and frontal areas (see **Figure 2C**). These areas are consistent with the some of the most important cortical regions that contribute to human processing of speech and language (Hickok and Poeppel, 2007). To ensure that the channels are in largely the same positions across participants, the probe sets were fitted on a specific cap based on the international 10-20 system (channel 7/29 corresponds to T7/T8 near the left/right primary auditory cortex). All participants wore the same cap. The vertex position and the nasion-vertex-inion midline were aligned across participants. To fit the channel positions on the cortical anatomy, the optodes and anatomical surface landmarks (nasion, vertex, inion, left and right ears) were registered using a 3D digitizer provided by the EGT-4000. In practice, it had shown difficult to appropriately register the landmarks in many of our participants (e.g., very small dislocations of digital sensors can cause greatly spurious head shape). Therefore, we used the most successful digitization result in one of the participants as the representative for channel positioning over the anatomical areas for all participants. Since a fixed cap was used, the standardized alignment procedure should *not* lead to large interindividual variability of channel positions with pronounced effects on the neural measurements.

Efforts were taken by the experimenters to maximize the good optode contacts with the scalp. With participants who had hair, a thin stick was used to help pull out the hair out of the way between the optodes and the scalp. General good contacts were ensured with waveforms having clear cardiac elements monitored by ETG-4000 in real-time. Formal tests started when better contacts could no longer be achieved after every effort was taken. Channels with poor signal quality were further detected and excluded for subsequent analyses (see *fNIRS data analyses*).

#### 2.4.2 Paradigms

The fNIRS experiments included an auditory and a visual test. The auditory test used speech and non-speech stimuli. The speech stimuli were ASL sentences spoken by the same male speaker as in the speech-in-noise tasks while non-speech stimuli were spectrally-rotated versions of the speech (Scott et al., 2000, 2009). The spectrally-rotated speech preserves some of the acoustic properties of the original speech, including similar wideband amplitude modulations, harmonic complexity and intonations, but they were highly unintelligible (Scott et al., 2000, 2009). This thus controlled for the auditory processing of acoustic properties that enabled us to study how neuroplasticity may be related to speech-specific factors such as intelligibility. All stimuli were presented via ER-3 earphones under an 8-talker babble noise with the overall sound level (sentence plus noise) calibrated at 75 dB as in the speech-in-noise tasks. The SNR was fixed across all sessions at the SRT obtained from the speech-in-noise task at T0 on a participant-by-participant basis. This ensured that speech stimuli were partly intelligible (∼50% word recognition at T0) which thus required similar listening efforts across participants and that neural responses to the speech/non-speech stimuli can be statistically compared across different sessions.

A block design was adopted in which participants sat in front of a computer screen with a grey background and a black cross in the middle for them to keep their eyes on and listened to 12 speech and 12 non-speech blocks presented in a randomized order (see **Figure 2B**). Each block consisted of 4 sentence trials. All sentences were ∼2 seconds long and each sentence plus noise was set to a fixed duration of 2.5 seconds that allowed the babble noise to start before sentence onset and extend after it ended. Another 2.5 seconds silent interval followed each sentence before the next during which participants were required to gently press a button (1, 2 or 3) to indicate how many key words they could recognize from the sentence. Each block thus lasted 20 seconds. Silent resting blocks were interleaved between the speech and non-speech blocks, each of which had a duration set randomly at 15, 17, 19 or 21 seconds. This was to reduce the possibility of participants being able to predict when the next speech/non-speech block would happen. The auditory test lasted for ∼15 minutes.

For the visual test, participants were exposed to a flashing radial chequerboard with black and white patches (the two colours reversed at the rate of 8 Hz, see Vafaee and Gjedde, 2000) on the computer screen against a grey background. Similar to the auditory test, a block design was used (see **Figure 2B**). There were 10 chequerboard blocks, each with a duration of 20 seconds. In addition to the chequerboard, a white cross appear in the middle of the screen and was set to change to red and then back to white (timings for the changes were set at random but occurred no earlier than 4 seconds after the block onset). To ensure participants’ engagement, they were asked to focus on the cross and gently press a button whenever the colour changed. Resting blocks were interleaved between stimulus blocks, each with a duration randomly set at 17, 20, or 23 seconds. The visual test lasted for ∼7 minutes.

A two to three minutes’ practice run was provided before formally starting each test so that participants were familiarized with the paradigms. Across the entire test period, participants were asked to restrain their body and head movements and consistently keep their eyes on the cross in the middle of the screen.

### 2.5 fNIRS data analyses

The signal processing procedure includes preprocessing, data processing of functional activations and connectivity, and statistical analyses. **Figure 3** shows the flow charts of this procedure.

**Figure 3.**
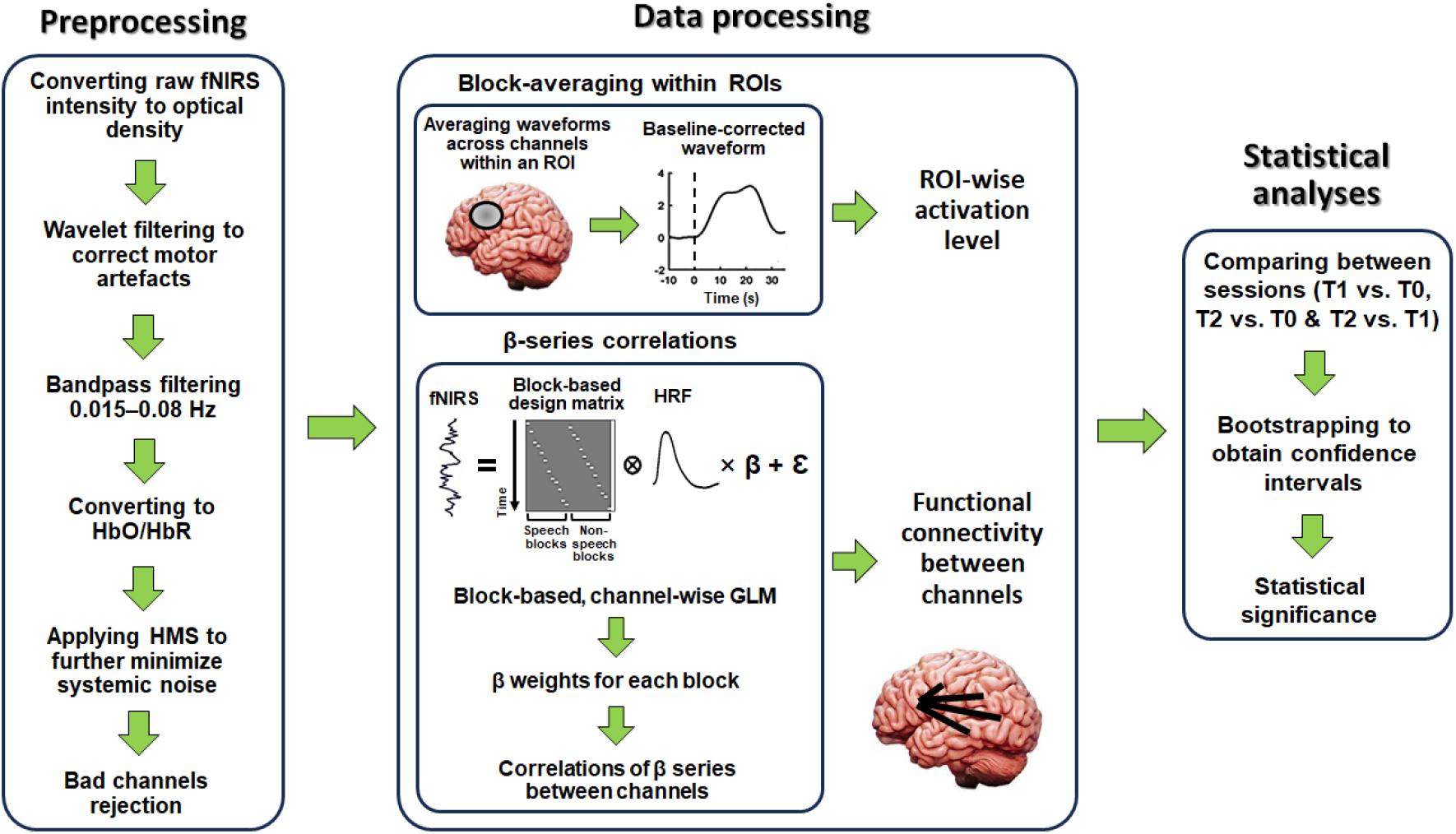
Flow charts for signal processing. The raw fNIRS data were first preprocessed. This included conversion fNIRS intensity to optical density, motor artefact correction (via wavelet filtering), bandpass filtering, conversion to HbO and HbR, and applying haemodynamic modality separation (HMS). Bad channels were finally detected via scalp coupling index (SCI) and were rejected for subsequent analyses. The preprocessed data were then used to measure functional activation levels and connectivity for each task (auditory and visual) during each session (T0, T1 and T2). Activation levels were measured via normalised response magnitudes by block-averaging within ROIs. Functional connectivity was measured by correlations of block-based beta-weight (through GLM) series between individual channels. Statistics were finally conducted using bootstrapping to obtain confidence intervals based on comparisons of activation levels and connectivity between different sessions for each task. Details of the entire procedure are described in the main text.

#### 2.5.1 Preprocessing

All signal processing and analyses of fNIRS were conducted using Matlab 2019b (Mathworks, USA) combining customized codes and the HOMER2 (Huppert et al., 2009) (homer-fnirs.org) and SPM-fNIRS toolbox (Tak et al., 2016) (www.nitrc.org/projects/spm_fnirs). We followed the signal processing procedure which was reported to result in high test-retest reliability of speech-evoked responses by fNIRS (Wiggins et al., 2016).

The raw fNIRS intensity signals were first converted to changes in optical density (via the HOMER2 function *hmrIntensity2OD*). Then motion artefacts were corrected using wavelet filtering (via the HOMER2 function *hmrMotionCorrectWavelet*). This removed wavelet coefficients lying more than 0.719 times the inter- quantile range below the first or above the third quartiles (Lawrence et al., 2018; Mushtaq et al., 2021). The optical density signals were then bandpass filtered between 0.015 and 0.08 Hz using a zero-phase 3^rd^-order Butterworth filter (hence covering the presentation frequency of ∼0.025 Hz in the block design) which attenuated the low-frequency drifts and changes in arterial blood pressure, respiratory and cardiac activities. The signals were then converted to changes in HbO and HbR concentrations via the modified Beer-Lambert Law (Huppert et al., 2009). The haemodynamic modality separation (HMS) algorithm (Yamada et al., 2012) was finally applied to further minimize the possible remaining systemic physiological noise and motion artefacts (e.g., slow head and body motions) (Wiggins et al., 2016). HMS is based on the fact that changes in HbO and HbR are negatively correlated in the functional responses but positively correlated in the motion and physiological noises. Accordingly, it returned separate estimates of the functional and noise components. We used the functional components for the changes in HbO as the final pre-processed measurements (due to the negative correlation with HbO, functional components for the changes in HbR were thus redundant after applying HMS, see Yamada et al., 2012).

As well as the pre-processing, channels with poor signal quality were detected despite the efforts to optimize optode contacts with the scalp. The scalp coupling index (SCI), which can effectively identify poor fNIRS signals in speech perception experiments (Pollonini et al., 2014; Mushtaq et al., 2019, 2021; Lawrence et al., 2021), were adopted. The signals with the two wavelengths were first bandpass filtered into 0.5–2.5 Hz that represents the cardiac elements captured by fNIRS and were correlated with each other. The higher correlation indicates better optode contacts. Following the criteria used in previous speech perception studies using fNIRS (Mushtaq et al., 2019, 2021; Lawrence et al., 2021), the worst 5% of channels (across all participants and sessions) were excluded for subsequent analyses. This threshold was set to ensure as many channels as possible (i.e., 95% of all channels) were preserved for statistical analyses (Mushtaq et al., 2019, 2021; Lawrence et al., 2021) especially when relative low number of participants (i.e., 10) were recruited in the current study.

#### 2.5.2 Data processing of functional activations and connectivity

The pre-processed fNIRS activations were analysed to measure: (1) functional activation levels; and (2) functional connectivity during both auditory and visual tests. We examined activation levels using block averaging across channels within several regions of interests (ROIs). This approach was employed because test-retest reliability in previous studies have shown that fNIRS activations are more reliably estimated when signals are averaged across small number of channels within a given ROI compared to when signals are analysed on a single-channel basis (Plichta et al., 2006; Schecklmann et al., 2008; Blasi et al., 2014; Wiggins et al., 2016). For the auditory tests, we focused on four ROIs of the bilateral auditory cortices (left: Channels 2, 3 and 7; right: Channels 24, 25 and 29), left inferior parietal lobule (Channels 11, 15, 16 and 20) and left frontal/prefrontal cortex (Channels 13, 17, 18 and 22). For the visual tests, we focused on two ROIs of the bilateral auditory cortices. Auditory cortices were chosen as we wanted to assess the functional neuroplasticity in auditory sensitivity in the auditory test and cross-modal maladaptation in the visual test. The other two ROIs (left inferior parietal lobule and frontal/prefrontal cortex) were chosen for the auditory test since they reflect higher-order speech and language processing dominant in the left hemisphere (Hickok and Poeppel, 2007). The left inferior parietal lobule is specifically associated with speech-in-noise perception (Alain et al., 2018) as well as semantic processing (Coslett and Schwartz, 2018), whilst left frontal/prefrontal cortex is associated with listening effort (Wild et al., 2012; Wijayasiri et al., 2017; Rovetti et al., 2019; Sherafati et al., 2022). The fNIRS waveforms were temporally averaged across channels within each given ROI for each trial. The averaged waveform was then baseline-corrected by subtracting the mean of the 10-second pre-stimulus period and normalized by dividing the pre-stimulus’ standard deviation (Balconi et al., 2015; Balconi and Vanutelli, 2016, 2017; Mutlu et al., 2020; Yorgancigil et al., 2022). The waveforms were then averaged across trials for each condition in each session. Because the haemodynamic responses peak at ∼5 seconds after the stimulus presentation, the response amplitude for a given condition was measured as the mean amplitude across the 5–25 seconds’ period (according to the 20 seconds block duration) after stimulus onset.

Functional connectivity was also quantified following the approach developed by Rissman et al. (2004) which measures correlations of beta-weight series across individual blocks (obtained via General Linear Model, GLM) between different channels. Specifically, design matrices were first created for the three experiment sessions (T0, T1 and T2) and for the auditory and visual tests, respectively. In each matrix, a boxcar regressor was created for every single block. The resting state was not included as a regressor based on the assumption that it did not actively trigger the haemodynamic responses and its activation level approximated to the global intercept. The canonical haemodynamic response function (HRF) was then convolved with the design matrix and the corresponding fNIRS signals were fitted using the convolved matrix via GLM (using the SPM-fNIRS toolbox) to obtain channel-wise beta weights. As such, a beta weight was obtained for every single block that reflected the level of activations of that block in each channel. This thus generated a beta-weight series for each condition (e.g., there were 12 blocks for the speech condition, hence giving a series of 12 beta values) for each channel. Pearson correlations of the beta-weight series were then calculated between individual channels (followed by Fisher-transform) as the values of connectivity between them. Such an approach has been successfully applied to quantify effective haemodynamic functional connectivity (Rissman et al., 2004; Ye et al., 2011; Gottlich et al., 2017; Antonucci et al., 2020; Pang et al., 2022).

### 2.6 Statistical analyses

Following acquirement of the behavioural (SRTs) and fNIRS data (activation levels and functional connectivity), statistically analyses were conducted to compare how these data changed between different experiment sessions (T1 vs T0, T2 vs T0 and T2 vs T1). Due to the relatively small number of participants, we applied bootstrapping instead of ANOVAs to avoid requirement for assumptions of specific data distributions (e.g., normality). Specifically, data were resampled with replacement in each replication and a bootstrap distribution was obtained after 10,000 replications. The confidence intervals were measured using the bias-corrected and accelerated (BCa) approach (using the Matlab function ‘bootci’) which corrected the confidence limits by accounting for deviations of the bootstrapped mean from the sample mean and skewness of the distributions (Efron, 1987; Efron and Tibshirani, 1994). An effect was considered as statistically significant if the value of zero fell outside the [1–α] (α as the significance level set at 0.05) confidence interval of a given distribution. For the SRTs and fNIRS activation levels in each ROI, α was set at 0.05/3 to correct for the number of sessions (i.e., 3). For the functional connectivity, α was set at 0.05/(946*3) to correct for the total number of connectivity between all 44 channels (i.e., 946) and the number of sessions (i.e., 3).

## 3 Results

### 3.1 Behavioural results

Behavioural speech-in-noise performances were measured as SRTs. We found significantly lower SRT (i.e., better speech-in-noise performance) at T2 than at T0, but no significant differences between T1 and T0 or between T2 and T1 (**Figure 4** and **Table 1**). This thus shows that speech-in-noise performance improved after retention (T2) but not immediately after training (T1).

**Figure 4.**
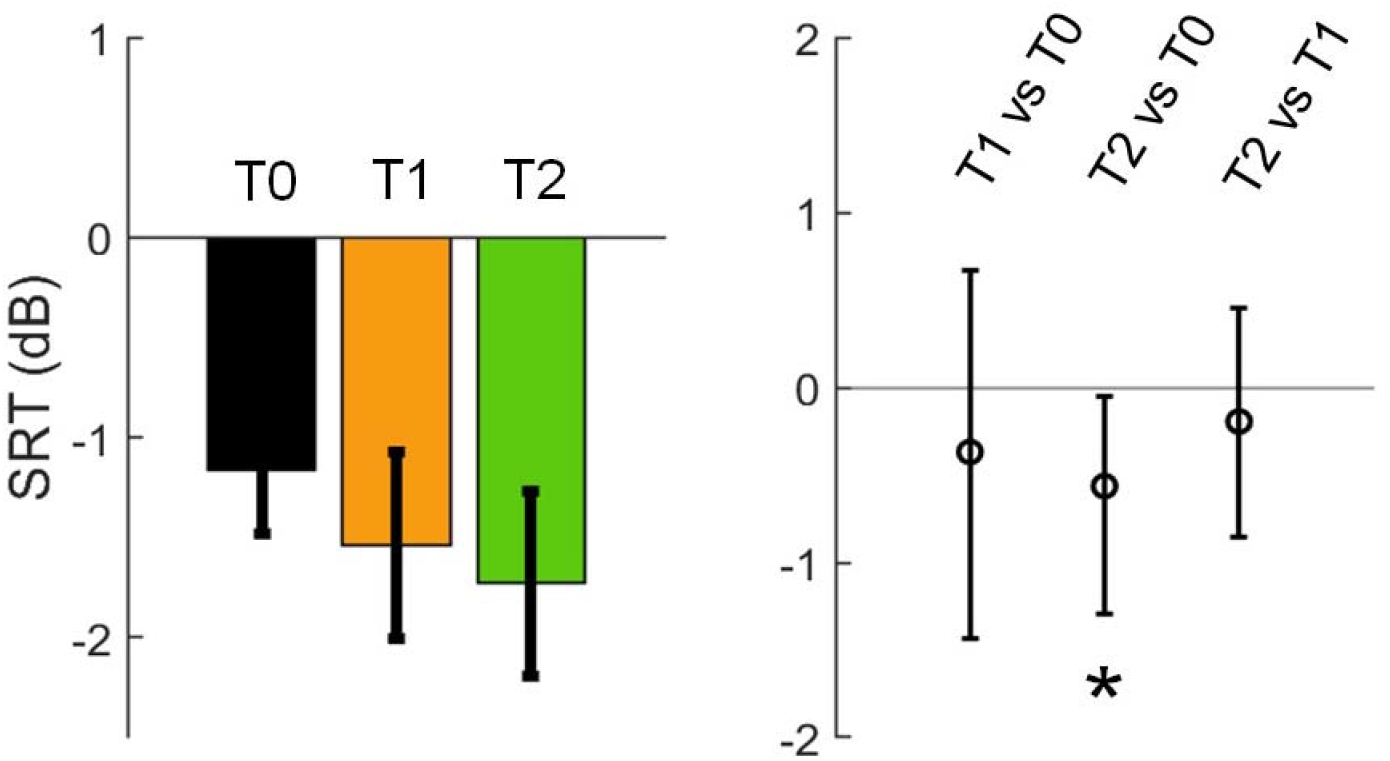
Speech-in-noise performances (SRT; lower SRT reflects better performance) across sessions. *Left panel*: SRTs at T0, T1 and T2. Error bars indicate standard errors of the means. *Right panel*: changes across sessions (T1 vs. T0, T2 vs. T0 and T2 vs. T1) with mean values indicated by circles in the middle and error bars indicating 95% confidence intervals (significance level α corrected at 0.05/3). The asterisk indicates significance where zero is outside the confidence interval.

**Table 1.** Statistical summary for the changes in SRT across sessions (T1 vs. T0, T2 vs. T0 and T2 vs. T1). Numbers in the brackets illustrate the 95% confidence intervals (significance level α corrected at 0.05/3). The asterisk indicates significance where zero is outside the confidence interval (in bold).

|  | T1 vs. T0 | T2 vs. T0 | T2 vs. T1 |
| --- | --- | --- | --- |
| SRT (dB) | [-1.455, 0.670] | **[-1.286, -0.048]\*** | [-0.877, 0.427] |

### 3.2 Neural results

#### 3.2.1 Auditory tests

Functional activation levels connectivity in response to auditory stimuli were compared between the three sessions. We conducted the comparisons separately for the speech and non-speech conditions, as well as for speech vs. non-speech.

For the activation levels, we found significantly changes in response magnitudes in the ROIs of left auditory cortex and left frontal/prefrontal cortex (see **Figure 5B** and **Table 2**). Specifically, response magnitudes were significantly increased at post-training (T1) than the baseline (T0) in the left auditory cortex for the non-speech condition and significant decreases in responses after retention (T2 vs. T0 and T2 vs. T1) for speech than non-speech. In the left frontal/prefrontal cortex, responses amplitudes were significantly reduced at post-training than baseline (T1 vs. T0 and T2 vs. T0) in the left frontal/prefrontal cortex for the speech but not the non-speech condition. In addition, such decreases were also significantly greater for speech than for non-speech. No significant differences were found between sessions in right auditory cortex or left inferior parietal lobule.

**Figure 5.**
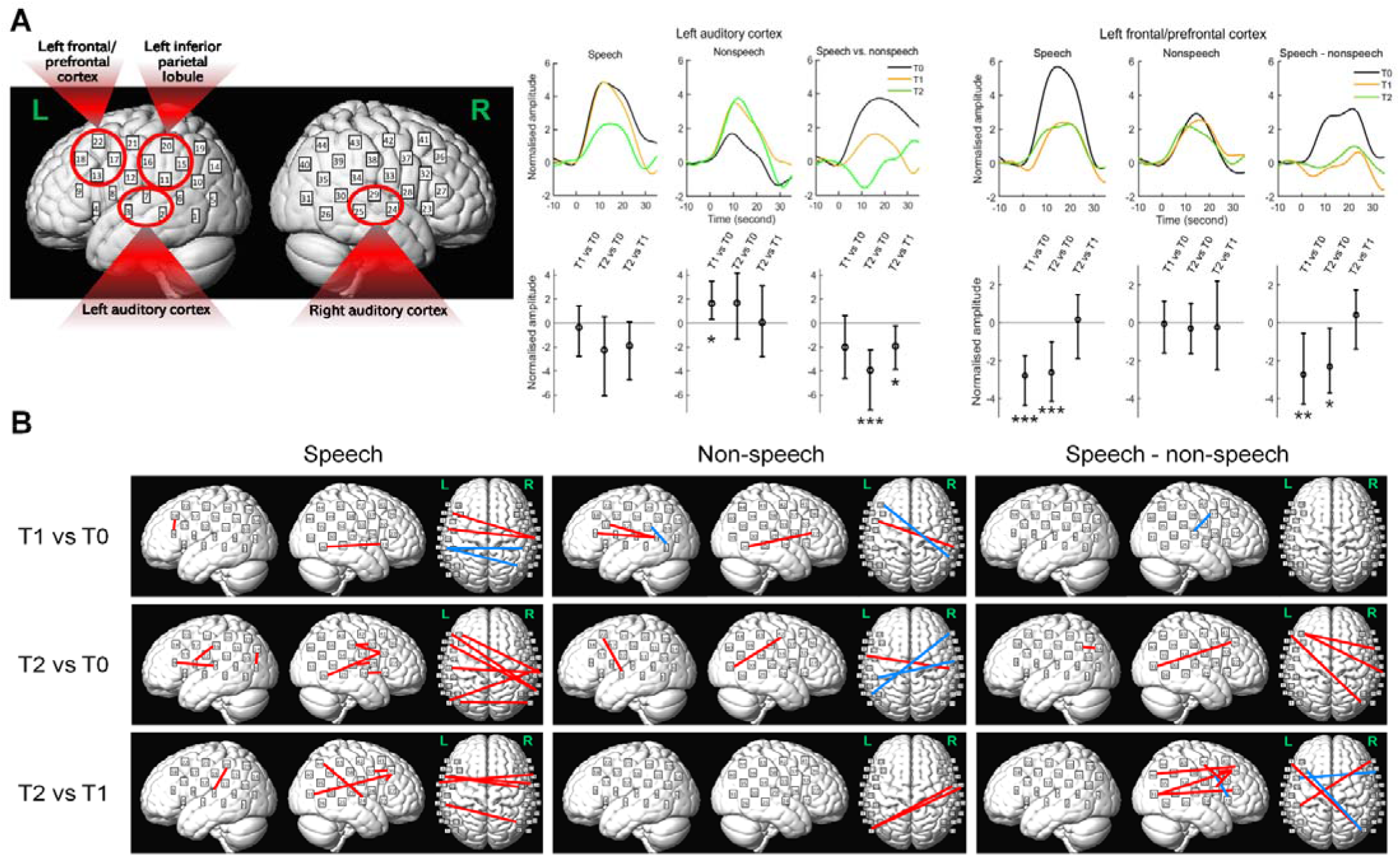
Changes in functional activation levels and connectivity during the auditory test across sessions (T1 vs. T0, T2 vs. T0 and T2 vs. T1) for the speech, non-speech and (speech - non-speech) conditions. See *Methods and Materials* for details of determining statistical significance. **(A)** *Left*: ROIs for calculating functional activation levels indicated by red circles. ROIs include the bilateral auditory cortices (left: Channels 2, 3 and 7; right: Channels 24, 25 and 29), left inferior parietal lobule (Channels 11, 15, 16 and 20) and left frontal/prefrontal cortices (Channels 13, 17, 18 and 22). *Right*: changes in response amplitude in the ROI of the left auditory cortex and left frontal/prefrontal cortex which showed significant changes (no significant changes between sessions were found for the right auditory cortex and left parietal lobule, hence patterns for these two ROIs were not shown here). The left auditory cortex had significant increases in amplitudes in T1 vs. T0 for non-speech and decreases after retention (T2 vs. T0 and T2 vs. T1). The left frontal/prefrontal cortex had significant decreases in amplitude after training for speech and speech vs. non-speech (T1 vs. T0 and T2 vs. T0). Averaged normalised amplitudes for all three sessions (upper panels) and changes across sessions with mean values indicated by circles in the middle and the error bars indicating 95% confidence intervals (significance level α corrected at 0.05/3, lower panels). Single, double, and triple asterisk(s) indicate that zeros are outside the 95%, 99% and 99.9% confidence intervals, respectively. **(B)** Changes in functional connectivity. In each panel, significant changes (α corrected at 0.05/(964*3)) in intra- (within left and right hemispheres respectively) and inter-hemispheric connectivity are shown respectively (from left to right). The red and blue lines indicate the enhancement/increases and decreases in connectivity, respectively, showing that major enhancement occurred for speech after retention (T2 vs. T0 and T2 vs. T1).

**Table 2.** Statistical summary for the changes in the activation levels across sessions (T1 vs. T0, T2 vs. T0 and T2 vs. T1) in specific regions of interest (ROI). Numbers in the brackets illustrate the 95% confidence intervals (significance level α corrected at 0.05/3). Single, double and triple asterisk(s) indicate that zeros are outside the 95%, 99% and 99.9% confidence intervals, respectively (in bold).

| Task | ROI | Condition | T1 vs. T0 | T2 vs. T0 | T2 vs. T1 |
| --- | --- | --- | --- | --- | --- |
| Auditory task | Left auditory cortex | Speech | [-2.846, 1.416] | [-5.848, 0.463] | [-4.919, 0.006] |
|  |  | Non-speech | **[0.284, 3.501]\*** | [-1.558, 4.231] | [-2.825, 3.000] |
|  |  | Speech vs. non-speech | [-4.590, 0.602] | **[-7.723, -2.235]\*\*\*** | **[-3.876, -0.217]\*** |
|  | Right auditory cortex | Speech | [-2.421, 2.318] | [-2.844, 1.064] | [-4.992, 1.467] |
|  |  | Non-speech | [-1.581, 3.462] | [-1.635, 2.562] | [-3.927, 2.835] |
|  |  | Speech vs. non-speech | [-2.417, 1.811] | [-2.985, 1.242] | [-3.108, 2.186] |
|  | Left inferior parietal lobule | Speech | [-3.296, 2.200] | [-5.494, 1.639] | [-3.182, 1.944] |
|  |  | Non-speech | [-1.782, 0.407] | [-2.153, 1.631] | [-1.236, 1.591] |
|  |  | Speech vs. non-speech | [-2.496, 2.823] | [-4.357, 1.580] | [-3.221, 2.554] |
|  | Left frontal/prefrontal cortex | Speech | **[-4.393, -1.766]\*\*\*** | **[-4.179, -1.058]\*\*\*** | [-1.969, 1.473] |
|  |  | Non-speech | [-1.705, 1.146] | [-1.593, 1.038] | [-2.481, 2.242] |
|  |  | Speech vs. non-speech | **[-4.261, -0.529]\*\*** | **[-3.732, -0.370]\*** | [-1.361, 1.709] |

For the functional connectivity, we found significant enhancements of connectivity for both speech and non-speech at T1 and T2 compared to T0 (as well as several decreases, see **Figure 5C**). Importantly, however, these enhancements were dominant in the speech condition after retention (i.e., T2). There were 14 pairs of channels for T2 vs. T0 and 9 pairs of channels for T2 vs. T1 for the speech condition as opposed to no more than 4 pairs of channels in any other comparison for speech/non-speech where significant enhancements were found. These enhancements include intra- and inter-hemispheric connectivity between auditory (channels 2, 3, 7, 23, 24 and 29) and non-auditory (parietal and frontal) channels. For speech vs. non-speech, significant enhancements were found between non-auditory channels (posterior temporal lobe, parietal and frontal lobes) for T2 vs T0 and T2 vs. T1 (**Figure 5C**). These changes in functional connectivity thus corresponded to the behavioural changes where speech-in-noise performances improved after retention (T2) but not immediately after training (T1).

#### 3.2.2 Visual tests

Same as the auditory test, brain activation levels (response amplitudes in ROIs) and functional connectivity for the visual tests were compared between sessions. For the activation levels, we did not find any significant differences in beta-weights in any channel or response amplitudes in either ROI (the left or right auditory cortex) between sessions.

For the functional connectivity, changes were mainly found in T1 where significant decreases in connectivity were found between 14 pairs of channels for T1 vs. T0, where only one pair was found for T2 vs. T0 (see **Figure 6**). Out of these 14 pairs for T1 vs. T0, only two pairs were those unrelated to auditory cortices (connectivity between channels 13 and 35 and between 5 and 36); the other 12 pairs were all between auditory cortices (10 pairs at channels 2, 3 and 7 on the left and 2 pairs at channel 24 on the right) and non-auditory regions in the parietal and frontal areas and temporo-parietal junctions. Therefore, the results show that brain connectivity between auditory cortices (especially the left auditory cortex) and higher-level non-auditory regions in response to the visual stimuli were significantly decreased immediately after training, but then such decreases vanished after retention.

**Figure 6.**
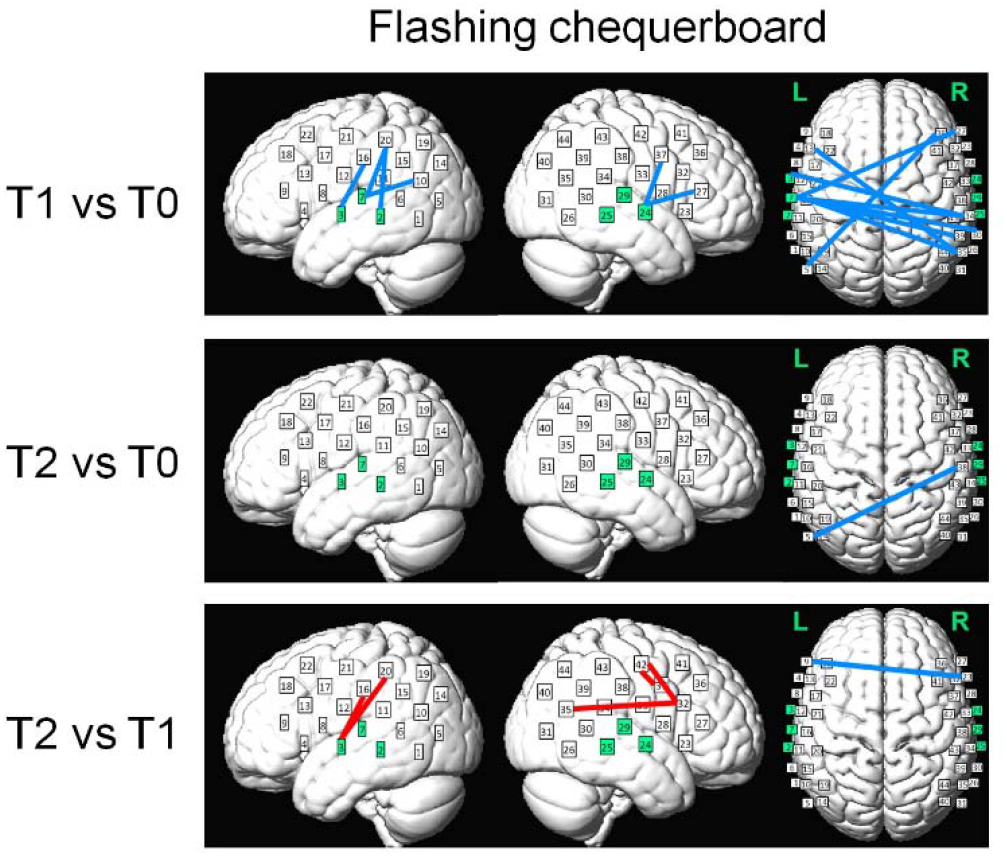
Changes in functional connectivity during the visual test across sessions (T1 vs. T0, T2 vs. T0 and T2 vs. T1). In each panel, significant changes (α corrected at 0.05/(964*3)) in intra- and inter-hemispheric connectivity are shown respectively (from left to right). The red and blue lines indicate the enhancement/increases and decreases in connectivity, respectively. Major changes were decreased connectivity between auditory and non-auditory cortices immediately after training (T1 vs. T0). Channels on the left and right auditory cortices (Channels 2, 3, 7, 24, 25 and 29) are highlighted as green.

## 4 Discussion

### 4.1 Neuroplasticity for speech-in-noise processing in noise older adults detected by fNIRS

Functional neuroimaging techniques, such as fMRI and PET, often face limitations in auditory research. These include loud scanning noise (e.g., fMRI) that requires careful design of paradigms in auditory experiments assuming that responses to the noise are the same across different experimental conditions (Hall et al., 1999, 2009; Blackman and Hall, 2011; Peelle, 2014). This could be tricky for hearing-impaired participants who often struggle with hearing in noisy backgrounds and when using speech stimuli who themselves are designed to be presented under noise. PET, on the other hand, does not have such caveat, but it is invasive requiring injection of radioactive isotopes, hence limiting its feasibility of repetitive use for longitudinal studies (Saliba et al., 2016; Basura et al., 2018; Harrison et al., 2021). Compared to fMRI/PET, fNIRS is non-invasive, acoustic silent/low noise and feasibly used longitudinally. In the current study, we used fNIRS to conduct a longitudinal study to examine auditory neuroplasticity in older adults. To our knowledge, there is the first study using fNIRS to examine neuroplasticity in terms of speech-in-noise perception. Most of our older adults (eight out of ten) had mild-to-moderate hearing loss, especially at high-frequencies (> 2 kHz), consistent with the real-life patterns of sensorineural hearing loss during normal ageing (Gopinath et al., 2009; Humes et al., 2010).

Older adults often face challenges in listening to speech under noisy environments (Humes, 1996), especially for those who have hearing loss (Souza and Turner, 1994; Barrenäs and Wikström, 2000; Humes, 2008) and speech-based training has been provided aiming to improve their speech-in-noise perception (Stropahl et al., 2020; Bieber and Gordon-Salant, 2021). Our results showed both behavioural and neural changes after training.

We showed significant behavioural improvements (i.e., speech-in-noise performances) after the retention period (T2), but not immediately after training (T1) compared to the pre-training baseline (T0) (**Figure 4**). This corresponded to enhancements in functional connectivity during the auditory tests. Significant enhancements in connectivity were predominantly observed for the speech condition at T2 (T2 vs. T0 and T2 vs. T1), but not T1 (T1 vs. T0) (**Figure 5B**). Such enhancements include greater intra- and inter-hemispheric connectivity between channels across bilateral temporal and parietal and frontal regions. This may indicate that changes in wide- spread functional connectivity could be potential indices for behavioural changes in speech-in-noise perception. This is also consistent with arguments that speech perception involves functioning of large-scale neural networks encompassing multiple wide-spread cortical regions that wire together rather than functioning of a single hub (Hickok and Poeppel, 2007). As indicated in our results, such networks whose enhancements were observed include not only lower-order auditory/temporal regions, but also higher-order non-auditory (parietal and frontal) regions. It has been reported that parietal cortices are involved with short-term phonological storage (Buchsbaum and D’Esposito, 2009), sensorimotor speech integration (Alho et al., 2014; Skipper et al., 2017) and semantic processing (Coslett and Schwartz, 2018), whilst frontal cortices are related to effortful listening (Wild et al., 2012; Wijayasiri et al., 2017), phonological working memory maintenance (Strand et al., 2008; Liebenthal et al., 2013) and syntactic processing (Grodzinsky et al., 2021) during speech perception. Also, the enhancements of inter-hemispheric connectivity indicate the potential importance of coordination and cooperation between the two hemispheres for speech-in-noise perception, which is a result, to our knowledge, that has not been reported previously.

We also found neural changes in the ROI of the left auditory cortex in the auditory tests that correspond to the behavioural changes. Intriguingly, we found significant decreases in functional activations in the left auditory cortex comparing speech with non-speech at T2 (T2 vs. T0 and T2 vs. T1) (**Figure 5A**). This is, however, *inconsistent* with our hypothesis predicting that auditory sensitivity, especially that to speech stimuli, should increase after training. A possible reason may be that the auditory stimuli consisted of not only target stimuli (speech/non-speech sentences), but also background noise (multi-talker babbles, see *2.4.2*). The decreased activations may thus be explained by suppression in neural responses to background noise in the left auditory cortex. It is noticeable that these significant decreases (speech vs. non-speech) were due to decreased responses in speech, while at the same time, increased responses to non-speech (see **Table 2**). We thus suggest that this can be interpreted as overall combined effects of neural suppression of background noise during speech listening (leading to decreased responses in the speech condition) and increases in general auditory sensitivity (leading to increased responses in the non-speech condition). This interpretation is consistent with previous studies showing that neural suppression of background noise could be more important than neural enhancement of target speech to achieve successful speech-in-noise perception in older adults with hearing loss (Peterson et al., 2017). Interestingly, these effects were significant only on the left, but not right, auditory cortex, further stressing the hemispheric specificity of speech processing (Hickok and Poeppel, 2007). To our knowledge, this is the very first finding of possible background suppression observed by haemodynamic responses to speech in noise according to longitudinal changes.

Furthermore, we also observed neural changes can occur *before* the significant changes in behavioural performances. Specifically, functional activation decreased in the left frontal/prefrontal cortex during the auditory tests at both T1 and T2 compared to T0, hence taking place before the behavioural improvements that only emerged at T2. These decreases occurred for the speech condition and were significantly greater for speech than non-speech (see **Figure 5A** and **Table 2**). This thus indicates that such effects were not merely driven by acoustics, but also higher-level speech-specific features like intelligibility. Previous research has demonstrated that activations in the left frontal/prefrontal regions reflect listening efforts during auditory and speech perception in populations with various hearing status, including young normal-hearing adults (Wild et al., 2012; Wijayasiri et al., 2017), older adults with normal hearing (Wong et al., 2009) and mild-to-moderate hearing loss (Rovetti et al., 2019), and cochlear implant patients who have severe hearing impairment (Sherafati et al., 2022). Therefore, this result demonstrated reduced listening effort during speech-in-noise perception even *before* the occurrence of behavioural improvement and such reduction persisted after the retention period.

We also observed significant decreases in functional connectivity between auditory cortices and non-auditory parietal and frontal regions during the visual (checkerboard flashing) test (T1 vs. T0), which also occurred *before* the significant behavioural changes. Previous studies have shown greater auditory cortical activities in hearing-impaired people when they process non-auditory (e.g., visual) stimuli possibly reflecting functional takeover of the auditory functions (Rouger et al., 2012; Campbell and Sharma, 2014; Chen et al., 2015; Dewey and Hartley, 2015; Corina et al., 2017) associated with worsened speech perception (Campbell and Sharma, 2014). The current result may thus reflect decreases in cross-modal takeover after training. Also, this result should be the first time to indicate the possible takeover effects reflected by functional connectivity between auditory and higher-order speech-related areas. This may also reflect greater suppression of activities in auditory-related areas during visual stimulations. However, such decreases did not persist after retention and thus did not correspond to the changes in speech-in-noise performances. We argue that this may be because older participants in the current study had either normal hearing or mild-to-moderate hearing loss, while the takeover effects shown in the previous studies were reported in those with severe hearing loss (Campbell and Sharma, 2014; Chen et al., 2015; Dewey and Hartley, 2015; Corina et al., 2017). It is thus possible that, with less impaired hearing, our participants may have lower potentials for cross-modal neuroplastic changes. Therefore, while these decreases were observed immediately after training, they may be harder to persist, especially when the training had stopped during the retention period. Nonetheless, we demonstrated these longitudinal changes in cross-modal activations in healthy older participants that have not been reported in previous studies, hence illustrating the promises of using fNIRS to study such changes in more hearing- vulnerable populations in the future.

Taken together, our results demonstrated the auditory neuroplasticity using fNIRS where longitudinal changes in brain functions in response to auditory and visual stimuli occurred along with changes in behavioural (i.e., speech-in-noise) performances. Specifically, we found that large-scale functional connectivity in response to speech in noise was enhanced corresponding to the behavioural improvements. We also found corresponding decreases in left auditory cortical responses to speech vs. non-speech, possibly reflecting neural suppression of background noise that contributes the behavioural improvements. Crucially, we also demonstrated that neural changes, i.e., decreased left frontal/prefrontal responses to speech (reflecting reduced listening efforts) and decreased visual-elicited connectivity between auditory cortices and higher-order speech-related non-auditory areas (reflecting reduced cross-modal takeover and/or greater cross-modal suppression), occurred *before* the emergence of behavioural improvements. These changes can thus be seen as neural precursors that would not be detected solely through behavioural measurements, hence indicating predictive/prognostic potentials for treatments of speech-in-noise perception in hearing-impaired populations.

### 4.2 Limitations and future research

The current finding that speech-in-noise performance was improved only after retention (T2) rather than immediately after training (T1) indicates that the training may have resulted in a longer/medium-term rather than an immediate behavioural effect. Alternatively, this may be due to learning effects of multiple experiment sessions. This would also apply to changes in neural activities observed here. Future studies including a control group without receipt of training would help to disentangle the training and learning effects. Nonetheless, an important goal of our study was to assess the promises of fNIRS to study auditory neuroplasticity alongside behavioural changes without much concerning about the exact driver of this plasticity. In this sense, it is less important to clarify the training and learning effects, whereas the speech-based training can be seen as a tool that helped facilitate the emergence of neuroplastic changes.

Another limitation was the small sample size. More participants would be recruited to have greater statistical power in the future and to allow for better estimation of how neural changes are associated with behavioural changes. We suggest that the bootstrapping approach applied here has mitigated this potential concern for sample size, but a larger sample size would be needed to validate our results by future studies. Also, future research would apply fNIRS in those who have more severe hearing impairment and/or those with hearing protheses (e.g., hearing aids and cochlear implants) to further prove the promises of fNIRS in wider hearing-vulnerable populations.

### 4.3 Conclusion

To our knowledge, the current study is the first to use the optical neuroimaging technique of fNIRS to test and observed longitudinal changes in auditory functions in older adults. fNIRS is a tool that has unique advantages to assess and monitor functional brain activities in hearing-vulnerable populations over other neuroimaging techniques like fMRI and PET. Here, we demonstrated evidence for detecting neuroplasticity for speech-in-noise perception using fNIRS. Novel findings of functional neural changes were illustrated along with behavioural changes in longitudinal experiments in older adults after speech-in-noise training. Corresponding to improvements in speech-in-noise performances, we observed increased functional connectivity across wide- spread speech- and language-related regions reflecting enhancement of inter-regional coordination/cooperation to process speech, as well as decreased left auditory cortical responses to speech in noise possibly reflecting neural suppression of background noise. More interestingly, neural changes not only occurred at the same time as behavioural improvements, but also emerged as neural precursors *before* these improvements took place. Specifically, listening effort to speech and cross-modal takeover was reduced (decreased activations in the left prefrontal/frontal cortex during speech listening and decreased connectivity between auditory cortices and higher-order non-auditory areas during exposure of visual stimuli, respectively) before speech-in-noise performances could be improved. To our knowledge, these novel findings have not been previously reported. The findings thus open up new opportunities for future studies to base on to further investigate neuro-markers for functional changes during speech processing in older adults. We also argue that the current study should lay the ground for evaluating auditory neuroplasticity in wider hearing-impaired populations in the future, such as those who wear hearing protheses (e.g., hearing-aid and cochlear implant users).

## Acknowledgement

All experiments were conducted at the neuroimaging laboratories based at UCL Department of Speech, Hearing and Phonetic Sciences. We thank the lab’s experimental officer Mr Andrew Clark for the technical assistance. We thank Dr Tim Green and Prof Stuart Rosen (UCL) for providing the speech-in-noise training program and the guidance for its application, and Prof Rosen for providing the Matlab scripts that create the spectrally-rotated speech. We also thank Dr Paola Pinti for useful advice on fNIRS signal processing. Parts of the Matlab scripts for signal preprocessing was provided by Dr Ian Wiggins (University of Nottingham).

## 5 Declarations

### Ethics approval

The current study was approved by the UCL Research Ethics Committee. All participants were consent and reimbursed for their participation.

### Data availability

data will be available upon request.

### Authors’ contributions

GM - conceptualization, funding acquisition, data collection, data analyses, original paper drafting, paper review and editing. ZJ – data collection, data analyses, paper review. XW – data collection, data analyses. IT – conceptualization, funding acquisition, paper review and editing, supervision. PH – conceptualization, funding acquisition, paper review and editing, supervision.

